# Expression of endogenous *Anopheles gambiae* microRNAs using an *Anopheles gambiae* densovirus (AgDNV) intronic expression system

**DOI:** 10.1101/2024.10.02.616337

**Authors:** Rebecca M. Johnson, Hillery C. Metz, Yasutsugu Suzuki, Kyle J. McLean, Jason L. Rasgon

## Abstract

*Anopheles gambiae* densovirus (AgDNV) is a highly species-specific parvovirus that reaches high titers in adult *Anopheles gambiae* mosquitoes with few transcriptomic effects and no significant fitness effects. Given these characteristics, AgDNV has been proposed as a viral vector for basic research and mosquito control. Previous work created an AgDNV co-expression system with a wild-type AgDNV helper plasmid and a transducing enhanced green fluorescent protein (EGFP) expressing plasmid that can be used to co-transfect cells to generate infectious recombinant transducing AgDNV virions. Generated virions infect the *An. gambiae* midgut, fat body, and ovaries, yet this viral vector system is limited in the size of transgenes that can be expressed due to capsid packaging limitations. Considering these size constraints, we created an artificial intron within the EGFP gene of the transducing construct that can express small pieces of genetic material such as microRNAs (miRNAs), microRNA sponges, or other small sequences. Placement of this intron in EGFP created a fluorescent reporter such that incorrect splicing produces a frameshift mutation in EGFP and an early stop codon whereas correct splicing results in normal EGFP expression and co-transcription of the intronic genetic cargo. A selection of miRNAs with predicted or demonstrated importance in mosquito immunity and reproduction with expression localized to the fat body or ovaries were chosen as intronic cargo, construct expression and splicing evaluated, and the impact of miRNA expression on putative miRNA targets was measured *in vitro* and *in vivo*. While the intron was correctly spliced in cells and mosquitoes, miRNA delivery resulted in inconsistent changes to miRNA and target gene transcript levels—possibly due to organ-specific miRNA expression or miRNA-target gene sequence mismatch. However, with optimization this viral vector and developed intron have potential as an expression tool within *An. gambiae* mosquitoes or cells.

## Introduction

*Anopheles gambiae* is the major vector of *Plasmodium falciparum* in sub-Saharan Africa, where most malaria cases occur [1]. While current malaria control efforts rely heavily on bite prevention via bed nets and mosquito reduction through insecticides, insecticide resistance is increasing [2]. Similarly, the efficacy of antimalarials used to treat disease in humans is threatened by drug-resistant parasites. As such, there is an increasing need for novel tools to better investigate *An. gambiae* biology as well as new methods for mosquito and pathogen control (such as the development of malaria-resistant mosquitoes capable of replacing susceptible populations) [3].

Although CRISPR-Cas9 editing has great promise, such experiments typically require specialized microinjection equipment, alterations to the mosquito genome, and the time-consuming establishment of mosquito lines. The introduction of genetic material without modifying the genome such as through genetically modified microbes offers an alternative approach that is useful for altering the expression of existing genes, introducing new genes, or, in the case of paratransgenesis, using a symbiont to express a transgene within the vector that acts against the pathogen. While the bacterium *Wolbachia* has been proposed for paratransgenesis and select strains have been successful at blocking pathogens in *Aedes aegypti*, *Wolbachia* has yet to be genetically modified and generating stable infections in *An. gambiae* has proven difficult [4–6]. Although there have been reports of *Wolbachia* infections in wild populations of *An. gambiae* in Africa, these findings need further verification and the impact of these infections on *P. falciparum* in *An. gambiae* are currently unclear [5,7–10]. Viruses also have potential for use in paratransgenesis, however, a limited number infect *An. gambiae* and few are ideal for genetic manipulation or for the introduction of new genes due to off-target effects, or danger of transmission to humans and resultant disease [11,12]. One of the only non-pathogenic insect-specific viruses known to infect *An. gambiae* was discovered in 2008 in *An. gambiae* Sua5B cells - the *An. gambiae* densovirus (AgDNV) [13]. AgDNV is closely related to other mosquito densoviruses including *Culex pipiens pallens* densoviruses (CppDNV) and *Aedes aegypti* densovirus (AaeDNV) and consists of a 4,139 nt ssDNA genome with terminal hairpins at each end that allow for viral genome packaging [13]. AgDNV is highly specific to *An. gambiae* and has poor infectivity even within closely related mosquitoes [14]. While some densoviruses in other mosquito species do cause mortality, AgDNV is not pathogenic to *An. gambiae* and increases in titer over the course of the adult lifespan while having little impact on *An. gambiae* fitness or gene expression [15–17]. These characteristics make AgDNV ideal for use as a late-life acting bioinsecticide or as a viral vector to express genes against the parasites themselves [16].

Previously a co-plasmid expression system was developed consisting of the unaltered AgDNV genome in a pBluescript cloning vector (pWTAgDNV) and a transducing pBluescript construct containing an *Actin5C* promoter, an *EGFP* reporter sequence, and a *SV40* termination sequence (pAcEGFP) [18]. (From here on, “p” designates the plasmid form of the construct whereas “v” designates the viral form; constructs lacking a label denote the general sequence.) As both plasmids in this co-expression system possess the AgDNV terminal hairpins that are crucial for genome packaging, both sequences get packaged into capsids produced by the wild-type construct. Virions produced by this co-expression system localize to mosquito tissues important for pathogen transmission and immunity such as the midgut, ovaries, and fat body when injected into adult mosquitoes [13,18]. Although AgDNV has potential as a viral vector, the small genome and capsid size limits the length of transgenes that densoviruses can express; for example, in AaeDNV, a size increase of 8% over that of the wild-type genome resulted in a 10% reduction in packaging efficacy [17,19]. While less deleterious, a shorter sequence can also negatively alter packaging efficacy [18]. These size constraints led us to modify the transducing construct of AgDNV to express small pieces of genetic material such as microRNAs or miRNA sponges that would increase the size of AcEGFP from 3,994 nt to closer to the 4,139 nt size of WT AgDNV but would not exceed the size of the WT genome [18].

MicroRNAs (miRNAs) are small, non-coding RNAs that act as post-transcriptional regulators of gene expression through the RNA interference (RNAi) pathway [20]. The RNAi pathway and miRNAs are highly conserved and in *An. gambiae*, over 163 miRNAs involved in a variety of processes including mosquito reproduction, immunity and development have been identified to date [21–27]. Endogenous miRNAs are coded in introns or intergenic sections of DNA and form short, hairpin-shaped secondary RNA structures following transcription [28]. After processing by Drosha, pre-miRNA hairpins are exported from the nucleus to the cytoplasm where they are cut further by Dicer into ∼22 nt duplexes [29]. One strand of each duplex is degraded, and the remaining, more stable strand forms the mature miRNA that is brought to target mRNA sequences by the RNA-induced silencing complex (RISC). The mature miRNA sequence binds to regions of mRNAs where there is sequence complementarity (often the 3’ UTR) and the degree of complementarity controls whether the mRNA transcript is cut and targeted for degradation or whether binding simply blocks translation [20,30]. Endogenous *An. gambiae* miRNAs or *in silico* designed miRNA sponges that are complementarity to mature miRNA sequences and “soak” up endogenous miRNAs are ideal for expression via AgDNV due to their small size [31,32]. Through the expression and endogenous processing of pre-miRNAs into mature miRNAs, levels of mature miRNAs can be enhanced whereas expression of miRNA sponges will lead to depletion of endogenous mature miRNAs.

As miRNAs are often encoded in introns or intergenic regions and we wanted to express small RNA cargo in a way that allowed for tracking of expression, we developed an artificial intron with a reporter phenotype within the EGFP gene of the transducing AgDNV construct (Fig 1A and 1B). To test this intron, we identified miRNAs for expression and manipulation that are putatively involved in mosquito functions such as immunity and egg development that are tied to organs that AgDNV is known to infect [33–36]. Selected miRNAs all had predicted or observed functions within *An. gambiae*, yet many have not been purposefully manipulated in *An. gambiae* and functional studies identifying specific mRNA transcript targets and downstream effects are lacking (Table 1) [35,37,38].

**Figure 1:**
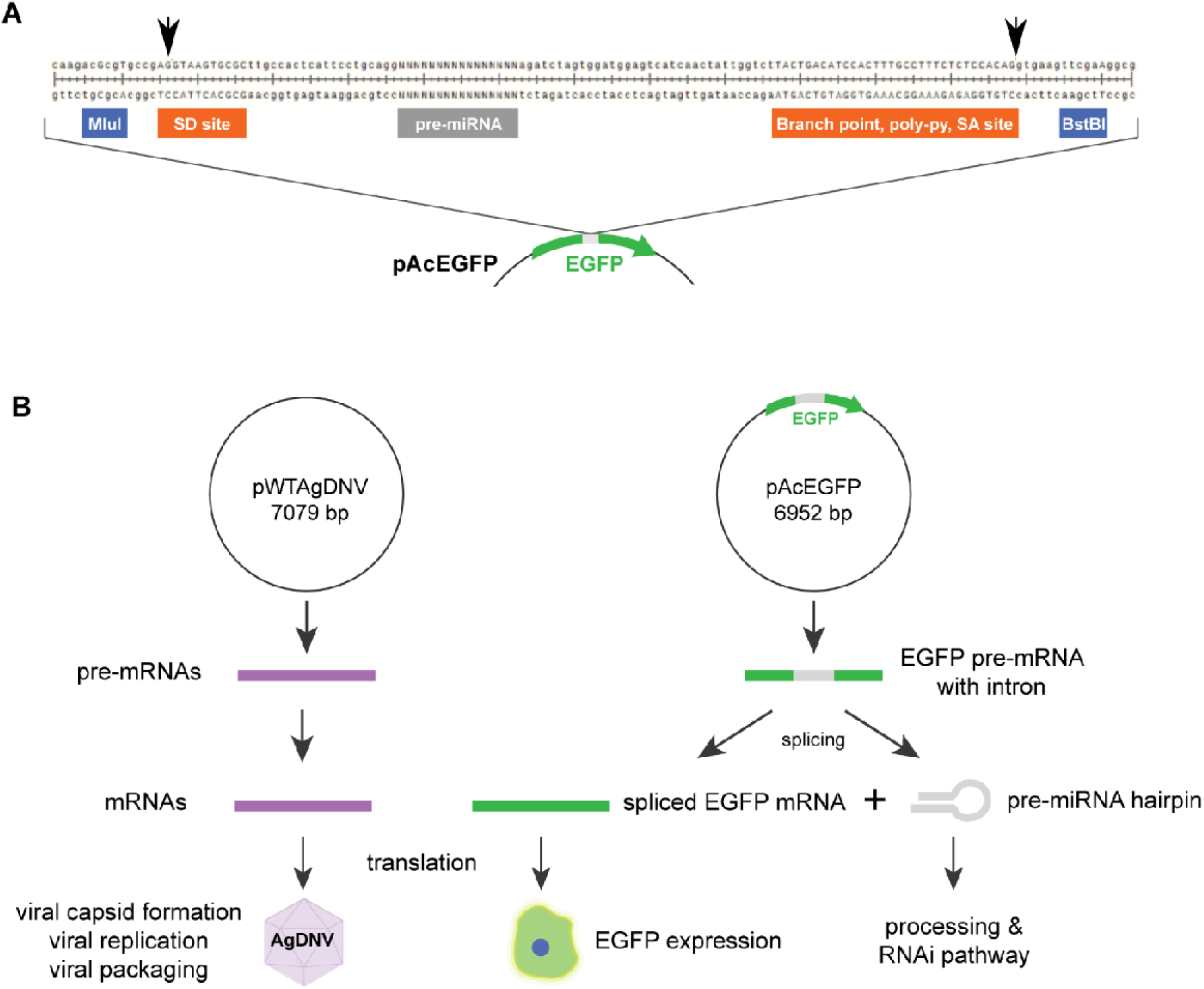
Intron layout and splicing scheme. **(A)** Splicing donor and acceptor sites within the EGFP gene flank the pre-miRNA region. *Mlu*I and *Bst*BI cut sites allow for swapping of g-block sequences containing *EGFP* bases removed during digestion, splice sites, and intronic cargo (pre-miRNA sequences, miRNA sponge, or *nonsense RNA*). Arrows mark cut sites within the splice donor and acceptor sequences. **(B)** During transcription, WT AgDNV transcripts are expressed from the WT construct (pWTAgDNV), while EGFP-encoding transcripts containing the intron are expressed by the transducing construct. Intronic splicing of the pre-mRNA (or miRNA sponge) transcript from the transducing construct results in the rejoining of EGFP-encoding transcript halves and the intronic cargo being processed via the RNAi pathway. Translation of WT DNV transcripts results in capsid formation whereas translation of EGFP mRNA results in EGFP expression if intronic splicing occurred correctly.

**Table 1:** Selected miRNAs and a miRNA sponge along with selected target gene transcripts. Many miRNA targets are predicted as *in vitro* and *in vivo* functional studies are lacking. Directionality, as noted through arrows, indicates an increase or decrease in target gene transcript levels following miRNA or sponge expression. Predicted target transcripts have no directionality as they have not yet been studied. The binding column notes whether this binding has been predicted or whether it has been observed in functional studies. Study species are noted to indicate which species these miRNAs and targets have been examined or predicted in.

| Construct | Target gene transcripts and directionality | Target gene functions | Target gene binding | Study species |
| --- | --- | --- | --- | --- |
| ***miR8*** | ↓ *Swim* | Egg development | Demonstrated | *Ae. aegypti*<br>*An. gambiae* |
| ***miR8SP*** | ↑ *Swim*<br>↓ *miR8* | Egg development | Demonstrated | *Ae. aegypti*<br>*An. gambiae* |
| ***miR34*** | *MISO* | Egg development | Predicted | *Ae. albopictus*<br>*Ae. aegypti*<br>*An. gambiae*<br>*An. stephensi* |
|  | *Cactus*<br>*Rel1* | Toll immune pathway regulation |  |  |
|  | *Caspar*<br>*Rel2* | IMD immune pathway regulation |  |  |
| ***miR305*** | *APL1C* | TEP1 regulation and *P. falciparum* immunity | Predicted | *An. gambiae*<br>*Ae. albopictus* |
| ***miR375*** | ↑ *Cactus*<br>↓ *Rel1* | Toll immune pathway regulation | Demonstrated | *Ae. aegypti* |
|  | *Caspar*<br>*Rel2* | IMD immune pathway regulation | Predicted |  |

The lack of validation of these selected miRNAs within *An. gambiae* proved challenging when assessing changes in miRNA or mRNA target levels, yet transcript splicing patterns indicate that the developed intron delivery system functioned as expected. This intron represents a new AgDNV viral vector expression strategy and may be useful for the expression of sequences including endogenous miRNAs, miRNA sponges, synthetic short interfering RNAs, or guide RNAs within genetically modified Cas9 mosquito lines.

## Methods

### Selection of miRNA targets

The first miRNA selected for this work, *miR8*, was highly upregulated in both the *Ae. aegypti* and *An. gambiae* fat body following blood feeding and targets the 3’ UTR region of secreted wingless-interacting molecule (*Swim*), a molecule involved with the Wnt/Wingless signaling pathway (Table 1) [39–42]. When *miR8* was depleted in *Ae. aegypti*, Swim levels remained high following blood feeding and egg development was inhibited [40]. Another miRNA, *miR34*, showed differential expression in several different mosquito species during pathogen infection including in *An. gambiae* where midgut was decreased following an infectious *Plasmodium berghei* (*P. berghei*) blood meal [21,33,43–45]. Specifically, *miR34* was predicted to bind to *Relish-like transcription factor 1*(*Rel1*) and *Caspar* transcripts, important factors in the Toll and IMD immune pathways respectively [36,45]. As *Caspar* is a negative regulator of the Relish like transcription factor 2 (Rel2) and *Cactus* is a negative regulator of *Rel1*, *Rel1* and *Rel2* transcript levels were also assessed in target gene qPCR reactions (Table S1). These transcripts as well as *mating induced stimulator of oogenesis (MISO)* were also predicted target genes of *miR34* via the now defunct miRNA-mRNA binding prediction webtool Insectar [46]. Previously, knockdown of *MISO* transcripts using RNAi resulted in reduced egg production, indicating a potential role for *miR34* in reproduction [47]. The third selected miRNA, *miR305*, was elevated in the ovaries and midgut of *An. gambiae* following blood feeding and higher in midguts following an infectious *Plasmodium*-containing blood meal [35,36]. Inhibition of *miR305* decreased the midgut microbiota and increased resistance to *P. falciparum* whereas enhancement of *miR305* increased *P. falciparum* infection levels and led to higher levels of midgut microbiota [36]. This miRNA was predicted to target the 3⍰ UTR of *APL1C* and as APL1 is part of a complex that stabilizes the immune factor thioester-containing protein 1 (TEP1), which binds to the surface of *Plasmodium* leading to parasite destruction, miR305 may impact *Plasmodium* infection [36]. Supporting this, *miR305* depletion in *An. gambiae* led to increased resistance to both *P. falciparum* and *Plasmodium berghei* infection and altered the levels of many immunity or anti-*Plasmodium* genes in mosquito midguts [42]. The final miRNA, *miR375*, was only detected in blood fed *Ae. aegypti* mosquitoes and was predicted to bind to the 5’ UTR of Toll pathway immune genes *Cactus* and *Rel1* [48]. Expression of a *miR375* mimic in *Ae. aegypti* mosquitoes or cells led to binding of the 5’ UTRs of *Cactus* and *Rel1* and the upregulation of *Cactus* and downregulation of *Rel1* [48]. Similar changes in target genes and an increase in Dengue virus type 2 titer was observed in *Ae. albopictus* Aag2 cell lines [48]. Although *miR375* has not been studied in *An. gambiae*, this miRNA has an identical sequence to *miR375* in *Ae. aegypti* and has also been predicted to target *An. gambiae Cactus* and *Rel1* along with other gene transcripts including *Caspar* and *Rel2* using Insectar [46].

### Plasmid preparation and production

Sure 2 supercompetent *E. coli* cells (Agilent Technologies, 200152) were transformed as per kit instructions (SOC media was substituted for NZY+ media) with pAcEGFP and pWTAgDNV plasmids [18]. Transformed colonies were plated on Luria broth agar plates with 100 µg/mL ampicillin and incubated at 37°C overnight. Colony PCR was used to verify transformation and selected colonies were grown in 5 mL of Luria broth in a 37°C shaker overnight and then preserved as glycerol stocks. Plasmids were produced by growing glycerol stocks in liquid culture as before, extracted using an Omega Bio-tek E.Z.N.A. Plasmid DNA Kit (D6942-02), and quantified using a NanoDrop ND-1000 spectrophotometer.

### Intron design

A potential splice acceptor site in WT AgDNV was identified at position 463 of the gene encoding the viral protein using the neural-network-based NetGene2 predictive splicing server which identifies transition sequences between introns and exons [49–51]. This sequence was converted from AG^ACGCAGACAG (with “^” indicating the predicted splicing site) into a splice donor site by replacing the intronic portion with the starting sequence of the second intron of *An. gambiae RPS17* such that the new sequence was AG^GTAGGCGCGC. This sequence was further modified by two base pairs to AG^GTA**A**G**T**GCGC to match the *An. gambiae U1* small nuclear RNA conserved region (Fig 1A) [52]. This U1 sequence (GTAAGT) represents the binding site for the *U1* small nuclear ribonucleoprotein which helps to form the spliceosome [52]. A splice acceptor with the sequence TACTGACATCCACTTTGCCTTTCTCTCCACAG was created to accompany this splice donor at position 464 of the gene encoding the viral protein by adding in the branch point, polypyrimidine tract, and intronic portion from the 3’ end of a chimeric human intron (last 32 nucleotides) preceding an immunoglobulin gene heavy chain variable region that is commonly found in commercial vectors such as in the pRL-CMV plasmid from Promega (Fig 1A) [53–55].

These developed *splice donor* and *splice acceptor*sites were placed within the EGFP gene of pAcEGFP at positions 334 and 337 respectively to create a reporter phenotype such that improper intronic splicing or a lack of splicing would result in a stop codon within the EGFP gene and correct splicing would result in EGFP expression (Fig 1B). Predicted splicing was examined at all steps using NetGene2 and the created *splice donor* and *splice acceptor* sequences both had a confidence scores of 1.0 indicating a high confidence in splicing [49].

### miRNA and sponge selection

For intronic miRNA expression, endogenous pre-miRNA sequences were inserted into the created intron so that upon splicing, the pre-miRNA hairpin would be co-transcriptionally processed alongside EGFP transcripts [21–23]. Selected pre-miRNA sequences for *An. gambiae miR8*, *miR34*, *miR305*, and *miR375* as well as a miRNA sponge against *miR8* (*miR8SP*) were added to this developed intron to test the co-expression system and intronic splicing mechanism. A random, *nonsense RNA* sequence (*NS*) was added to the intron as a control. These miRNAs and miRNA sponge were chosen based on known or predicted effects on genes involved with immunity, pathogen defense, or reproduction in *An. gambiae*, *Ae. aegypti*, or relevant mosquito cell lines (Table 1). To test intron functionality and demonstrate that splicing is sequence-dependent, altered *splice donor* and *splice acceptor* site sequences were developed using site-directed mutagenesis of the pAcEGFPmiR8 plasmid [49]. When the *splice donor* site was changed by a single nucleotide (in bold) from AG**G**TAAGTGCGC to AG**A**TAAGTGCGC, NetGene2 no longer identified this as a splice donor site. Similarly, when the *splice acceptor*site was changed by one nucleotide (in bold) from TACTGACATCCACTTTGCCTTTCTCTCCACA**G** to TACTGACATCCACTTTGCCTTTCTCTCCACA**T**, this site was no longer predicted to be a splice acceptor.

### Cloning and intronic cargo

*Mlu*I and *Bst*BI sites were introduced into the EGFP-encoding gene of pAcEGFP using site-directed mutagenesis to create synonymous mutations. A *Mlu*I site was created by altering position 327 of *EGFP* from a C to a G and position 330 from a C to a T. A *Bst*BI site was created in *EGFP* by switching position 348 from G to A. Endogenous *An. gambiae* pre-miRNA sequences from miRbase were converted to DNA and used to order g-blocks from Integrated DNA Technologies (IDT) [21]. The *mir8SP* sequence contained ten repeated blocks of the reverse complement of mature *An. gambiae miR8*. Each block was separated by four spacer nucleotides and the entire sponge sequence was placed within the intron as with pre-miRNA sequences. A *nonsense RNA* (*NS*) was created using a random sequence with no matches to the *An. gambiae* genome or transcriptome when searched using Basic Local Alignment Search Tool (BLAST). Each pre-miRNA, miRNA sponge, or nonsense RNA was coded on IDT g-blocks synthesized with flanking *Mlu*I and *Bst*BI sites, *EGFP* segments to replace those removed during digestion, and the *splice donor* and *splice acceptor* sites (Table S2; Fig 1A). G-blocks were subcloned into pJet using a CloneJet PCR Cloning Kit (ThermoFisher Scientific, K1231) and later digested using *Mlu*I and *Bst*BI. These inserts were ligated into pAcEGFP that had been digested with *Mlu*I and *Bst*BI and resulting plasmid sequences were verified.

### Cell culture and transfections

Sua5B and Moss55 *An. gambiae* cells were grown in 25 cc plug cap flasks at 28°C and passaged once per week at a 1:5 dilution with Schneider’s *Drosophila* media with 10% FBS v/v. For transfections, cells were quantified using a hemocytometer and 6 × 10^6^ cells were added to each well of a 6-well plate along with 3 mL of complete media and incubated overnight. Cells were transfected at ∼70-80% confluence with a 1:2 ratio of pWTAgDNV to transducing plasmid with 830 ng pWTAgDNV and 1660 ng transducing plasmid per well using a Lipofectamine LTX with Plus Reagent kit (ThermoFisher Scientific, 15338030). Briefly, plasmids were added to a mix of 500 µl OptiMem media with 3 µl Plus reagent and incubated at room temperature for 10 minutes. Then, 5 µl lipofectamine was added and tubes were incubated at room temperature for 25 minutes before transfecting each well with 500 µl of this mixture. Transducing plasmids were pAcEGFPmiR8, pAcEGFPmiR8SP, pAcEGFPmiR34, pAcEGFPmiR305, pAcEGFPmiR375, pAcEGFPNS, pAcEGFPSA, and pAcEGFPSD. Cells were incubated and imaged at 3 days post-transfection. RNA for splicing validation was also gathered 3 days post-transfection. Preliminary *in vitro* miRNA and target gene expression experiments harvested RNA 5 days post transfection. For Sua5B *in vitro* miRNA and target gene expression, cells transfected with pWTAgDNV and pAcEGFPNS served as controls whereas in Moss55 *in vitro* miRNA and target gene expression experiments, cells transfected with pWTAgDNV alone served as a control.

### Viral production and quantification

To produce virus particles for mosquito infections, Moss55 cells were transfected with selected transducing and helper viruses as described and virions extracted 3 days post-transfection by removing the media, washing cells with 1X PBS, and suspending cells in 1 mL 1X PBS. Cells were lysed using three cycles of freeze-thawing and centrifuged at 5,000 rpm for 5 minutes to pellet debris. The virus-containing supernatant was collected and plasmid DNA and free viral genomes were removed using an Ambion TURBO DNA-*free* kit (AM1907). DNA was extracted using an Omega Bio-tek E.Z.N.A Tissue DNA kit (D3396-02) and viral genome equivalents were determined using standard curves created using AgDNV-coding plasmids with a single copy of each gene-of-interest. Samples and standards were run using PerfeCTa SYBR Green FastMix (Quantabio, 95072-012) on a Qiagen Rotor-Gene Q at 95°C for 2 minutes followed by 40 cycles of 95°C for 10 seconds, 60°C for 40 seconds, and 72°C for 30 seconds. Runs were finished with a melt step using a ramp of 55°C to 99°C rising by one degree each step. WT AgDNV was quantified using primers against *AgDNV nonstructural gene 1* (NS-RT-IIIF: CATTCGATCACGGAGACCAC, NS-RT-IIIR: GCGCTTGTCGCACTAAGAAAC) and a standard curve of pWTAgDNV. Selected transducing viruses (vAcEGFPmiR8, vAcEGFPmiR8SP, vAcEGFPmiR34, vAcEGFPmiR305, vAcEGFPmiR375, vAcEGFPNS, vAcEGFPSA, and vAcEGFPSD) were quantified using primers against *EGFP* (GFP-RT-II-F497: TCAAGATCCGCCACAACATC, GFP-RT-II-R644: TTCTCGTTGGGGTCTTTGCT) and a standard curve of pAcEGFP. Each production of virus consisted of a mixture of vWTAcEGFP and a transducing virus.

### Mosquito injections

Female *An. gambiae* mosquitoes (Keele strain) that were 3 days post-emergence were injected intrathoracically with 200 nl densovirus mixture containing both wild-type vWTAgDNV and transducing virus (either vAcEGFPmiR8, vAcEGFPmiR8SP, vAcEGFPmiR34, vAcEGFPmiR305, vAcEGFPmiR375, vAcEGFPNS, vAcEGFPSA, or vAcEGFPSD) using a Drummond Scientific Nanoject III (3-000-207) and Drummond Scientific 10 µl microcapillary tubes (3-000-210-G) pulled using a Sutter Instrument Co. Model P-2000 (Heat=400, Fil=4, Vel=40, Del=140 Pul=140). Three biological replicates in mosquitoes were completed. For replicate 1, all vWTAgDNV and transducing virus combinations were used and mosquitoes were harvested at 10 days post-injection. For replicates 2 and 3, mosquitoes were only injected with vWTAgDNV and vAcEGFPmiR34, vAcEGFPmiR375, or the control vAcEGFPNS and harvested 10 days post-injection. For each replicate, mosquitoes were injected with ∼10^6^-10^7^ transducing virus particles and 10^6^-10^8^ WT DNV particles (Table S3). Mosquito treatment groups were kept in separate cardboard cup cages with 10% sugar solution w/v *ad libitum* until RNA extraction or imaging. RNA was harvested and tested from three biological replicates.

### RNA extractions and cDNA production

For both *in vitro* and *in vivo* experiments, RNA was extracted using an Omega Bio-tek MicroElute Total RNA Kit (R6831-02). For *in vitro* experiments, RNA was extracted at 3 days post-transfection for intronic splicing assessments or 5 days post-transfection for miRNA and target gene quantification. For *in vivo* experiments, mosquitoes were individually homogenized 10 days post-injection in lysis buffer using zinc-plated steel BB pellets (Daisy 0.177 cal or 4.5 mm) and a Qiagen TissueLyser II with a lysis program lasting 2 minutes with a frequency of 30 times/second. Following homogenization, RNA was extracted and DNase treated either on the column using an Omega Bio-tek RNase-free DNase Set I kit (E1091) or following RNA extraction using an Ambion DNA-*free* DNA Removal Kit (AM1906). For target gene quantification or assessment of intronic splicing, cDNA was synthesized using a Quantabio qScript cDNA synthesis kit (95047-500) whereas for miRNA quantification, samples were converted to cDNA using the HighSpec option in the Qiagen miScript II RT kit (218161) and diluted 1:10.

#### Intronic splicing, miRNA expression, and target gene quantification

*In vitro* and *in vivo* intronic splicing was assessed using primers spanning the intronic region (GFP-COLPCRF: CTGACCTACGGCGTGCAGTGC, RGFP-COLPCRR: CGGCCATGATATAGACGTTGTGGC). PCR products were run on 2% agarose gels and imaged using a UVP GelDoc-It transilluminator. Spliced transcripts resulted in a product of 274 bp whereas PCR reactions using DNA plasmid controls or unspliced transcripts produced variably sized amplicons depending on insert size with most being ∼480 bp.

*In vitro* experiments and replicate 1 *in vivo* target gene qPCR reactions were run on a Qiagen Rotor-Gene Q using PerfeCTa SYBR Green FastMix (Quantabio, 95072-012) with conditions of 95°C for 2 minutes, 40 cycles of 95°C for 10 seconds, 60°C for 40 seconds, and 72°C for 30 seconds and a melt curve with a ramp from 55°C to 99°C with 1-degree change per step. *In vivo* replicates 2 and 3 used Applied Biosystems PowerUp SYBR Green Master Mix (A25724) and an Applied Biosystems 7900HTFast Real-Time PCR System with 384-well plates and conditions of 50°C for 2 minutes, 95°C for 10 minutes, and 40 cycles of 95°C for 15 seconds, and 60°C for 1 minute. A dissociation step of 95°C for 15 seconds, 60°C for 15 seconds, and 95°C for 15 seconds ended each run. Primers for *An. gambiae Swim* cDNA were developed during this study whereas others came from published studies (Table S1) [47,56–59].

Reactions to quantify miRNAs used Qiagen miScript SYBR Green PCR kits (218075) and a Qiagen Rotor-Gene Q with a universal reverse primer and forward primers consisting of the sequence of each mature miRNA (Table S4) [39]. *An. gambiae U6* levels served as a reference with which to compare miRNA levels.. Conditions for miRNA qPCR reactions were 95°C for 15 minutes followed by 40 cycles of 95°C for 15 seconds, 60°C (for all miRNAs during cell culture replicates as well as for *in vivo miR34*) or 55°C (all *U6* reactions and *in vivo* miR375) for 60 seconds, and 72°C for 20 seconds. All reactions ended with a melt curve consisting of a ramp from 55°C to 99°C that increased one degree per step.

#### Data analysis

All qPCR data was analyzed using the delta-delta Ct method to calculate the fold change in expression relative to reference genes (*S7* for mRNA transcripts or *U6* for mature miRNA quantification unless otherwise noted). The fold change expression data was log2 transformed and a D’Agostino-Pearson omnibus K2 test was used to assess normality in Graphpad Prism 9. If both the control and experimental groups passed the normality test, a parametric unpaired t-test assuming equal standard deviations was used to measure statistical significance. If either or both groups failed the D’Agostino-Pearson normality test, a nonparametric Mann-Whitney test was used to compare ranks and to assess significance. Significant p values (<0.05) were reported on graphs. All graphs report fold change expression using a log2 scale. The mean and standard error of the mean was reported for groups analyzed using an unpaired t-test whereas median and 95% confidence intervals were shown for groups compared using a nonparametric Mann-Whitney test.

## Results

When *An. gambiae* Sua5B or Moss55 cells were co-transfected with pWTAgDNV and the original transducing plasmid pAcEGFP that lacked the created intron, strong EGFP expression was observed (AcEGFP, Fig 2A, B). Sua5B and Moss55 cells that were co-transfected with pWTAgDNV and transducing plasmid pAcEGFPmiR8, pAcEGFPmiR8SP, pAcEGFPmiR34, pAcEGFPmiR305, or pAcEGFPmiR375 had visible EGFP expression indicative of intronic splicing 3 days post-transfection (miR8, miR8SP, miR 34, miR305, and miR375, Fig 2A, B). EGFP expression was also present when cells were transfected with pWTAgDNV and the *nonsense-RNA*-encoding transducing plasmid pAcEGFPNS (NS RNA, Fig 2A, B). When *splice donor* or *splice acceptor*sites were mutated, EGFP expression was not detectable in Sua5B cells 3 days post co-transfection with pWTAgDNV and pAcEGFPSD or pAcEGFPSA, indicating that splicing was dependent on splice site sequences (SD mutant and SA mutant, Fig 2A). In *An. gambiae* Moss55 cells, similar expression patterns were observed (Fig 2B). EGFP signals were generally weaker in Moss55 compared to Sua5B cells, however results were consistent from both cell lines and indicate that intronic splicing is induced in a sequence specific manner for a wide variety of pre-miRNAs, miRNA sponges, and small RNAs in *An. gambiae* cell lines of varied lineage.

**Figure 2:**
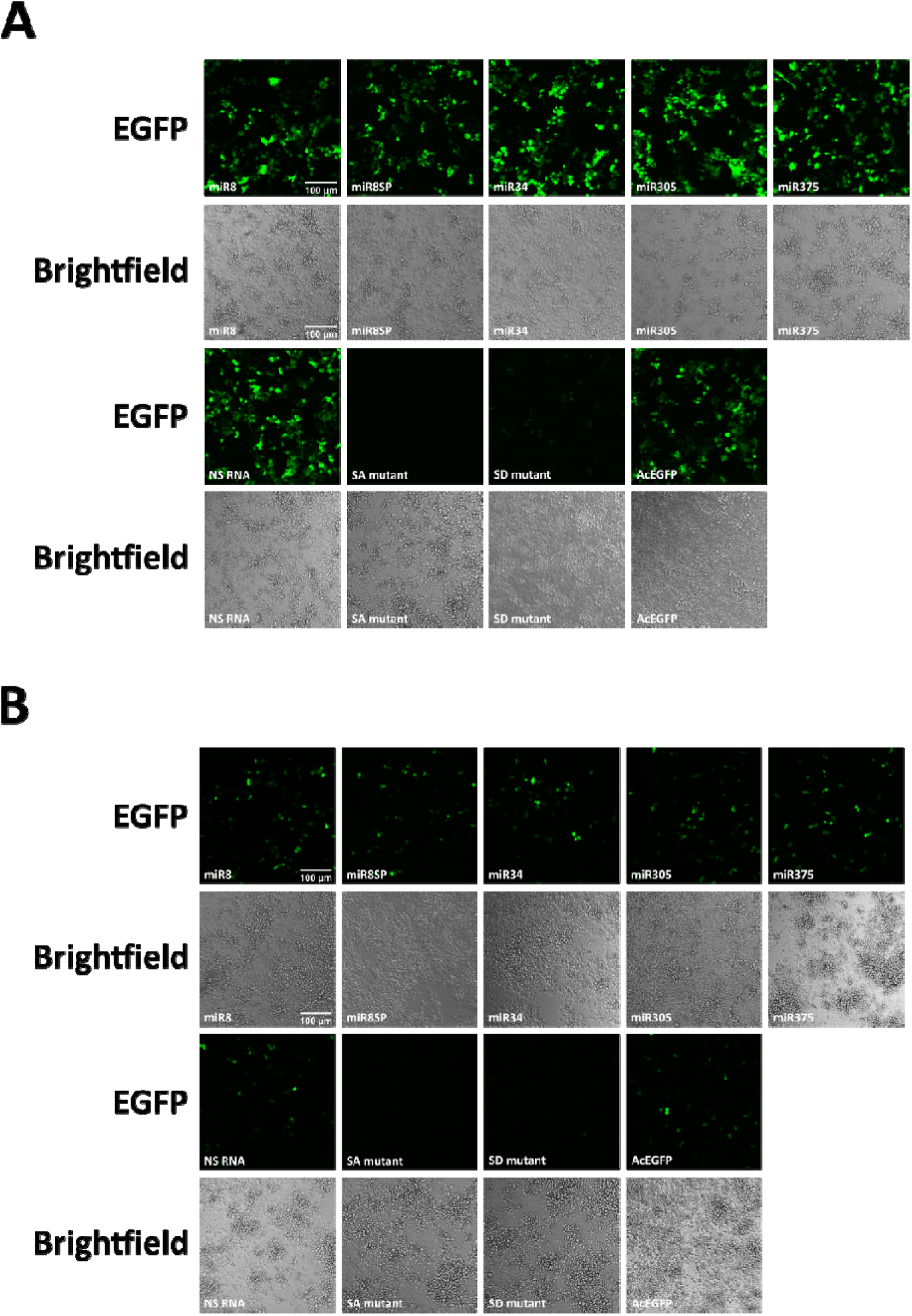
EGFP expression in Sua5B and Moss55 cells 3 days after co-transfection with pWTAgDNV and selected transducing plasmids. **A)** EGFP expression in co-transfected Sua5B cells. **B)** EGFP expression in co-transfected Moss55 cells. Panels are labeled with the transducing construct that was co-transfected with pWTAgDNV. Scale=100 um.

*In vitro* intronic splicing was further validated using primers spanning the intronic insert. A 274 bp PCR product consistent with splicing was observed in Sua5B cells 3 days post co-transfection with pWTAgDNV and transducing constructs pAcEGFPmiR8, pAcEGFPmiR8SP, pAcEGFPmiR34, pAcEGFPmiR305, pAcEGFPmiR375, and pAcEGFPNS (cDNA wells 1-6, Fig 3A). Unspliced plasmid DNA samples had PCR product sizes dependent on intronic length. This was ∼480 bp for most constructs except the *miR8SP*-expressing plasmid which had a larger insert and an amplicon of 572 bp (plasmid wells, Fig 3A). Both cDNA and plasmid versions of *EGFP* lack the intron sequence and have the same PCR product size of 274 bp (cDNA well 10, plasmid well 10, Fig 3A). Sua5B cells co-transfected with pWTAgDNV and constructs containing mutated *splice acceptor* or *splice donor* sequences (pAcEGFPSA or pAcEGFPSD) exhibited some level of intron splicing despite absent or greatly reduced visible EGFP expression (cDNA wells 7 and 8, Fig 3A; SD mutant and SA mutant, Fig 2A). This indicates that some transcripts are spliced despite the lack of predicted splicing via NetGene2 but that this splicing may be incomplete or in a location that causes a disruption in EGFP expression due to a stop codon. Faint ∼480 bp bands indicating the presence of some unspliced transcript were also observed in PCR reactions using cDNA from cells co-transfected with pWTAgDNV and pAcEGFPNS (cDNA well 6, Fig 3A). This points to some level of splicing disruption in these constructs yet, given the visible EGFP expression in cells co-transfected with pWTAgDNV and pAcEGFPNS, this may be explained by splicing intermediates (NS RNA, Fig 4A).

**Figure 3:**
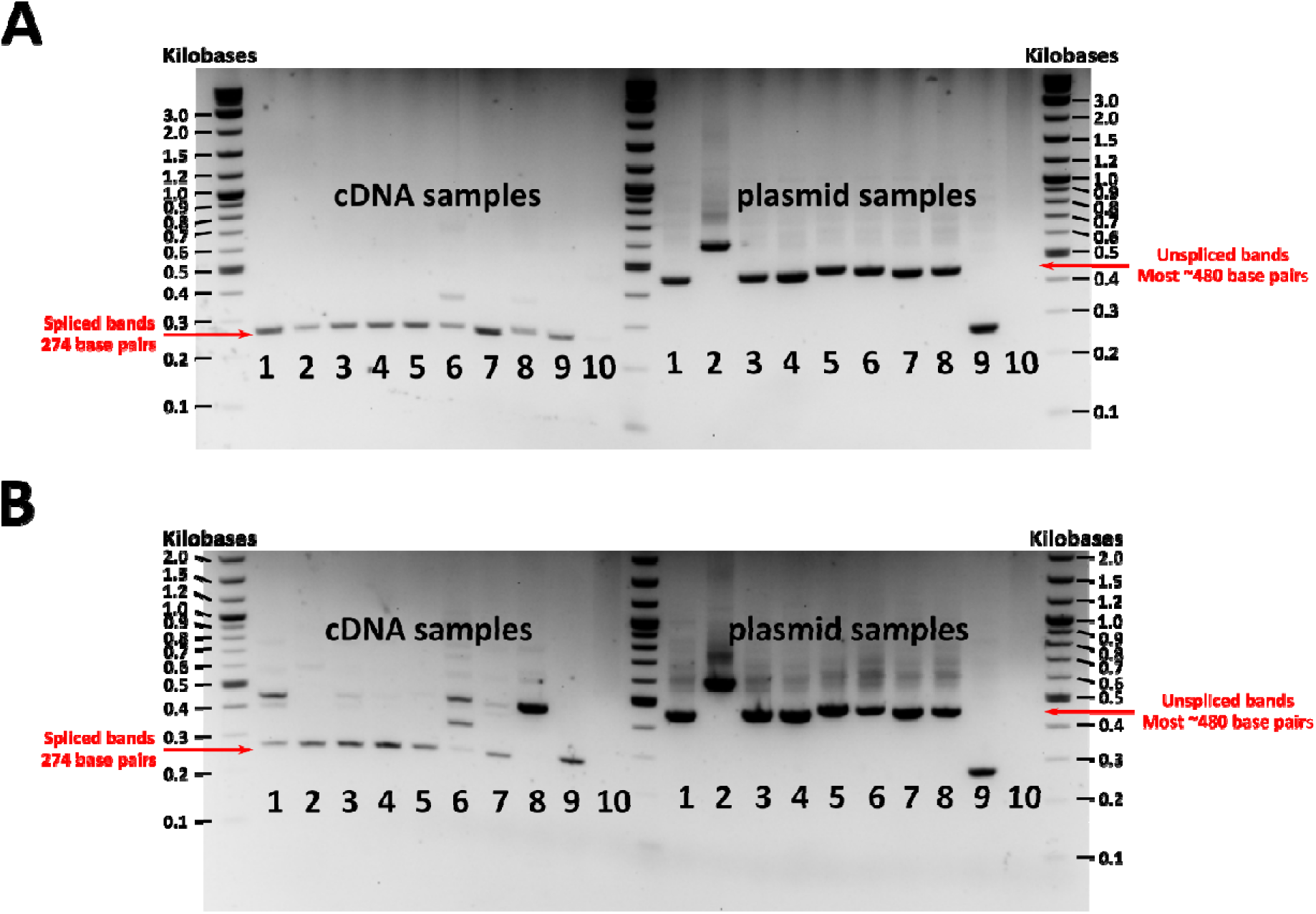
Intronic splicing in Sua5B and Moss55 cells 3 days post co-transfection alongside unspliced DNA plasmid control samples. **A)** Intronic splicing and plasmid controls in co-transfected Sua5B cells. **B)** Intronic splicing and plasmid controls in co-transfected Sua5B cells. In both **A** and **B**, Matching numbers indicate cDNA and plasmid versions of the same construct. Samples are as follows: 1. *miR8*, 2. *miR8SP*, 3. *miR34*, 4. *miR305*, 5. *miR375*, 6. *Nonsense RNA*, 7. *SA* mutant, 8. *SD* mutant, 9. *EGFP* lacking the intron, 10. No template control. Splicing of the intron in cDNA samples resulted in a PCR product of 274 bp whereas a lack of splicing as observed in plasmid controls on the right side of the gel resulted in bands of ∼480 bp for most constructs and 572 bp for miR8SP in well 2.

**Figure 4:**
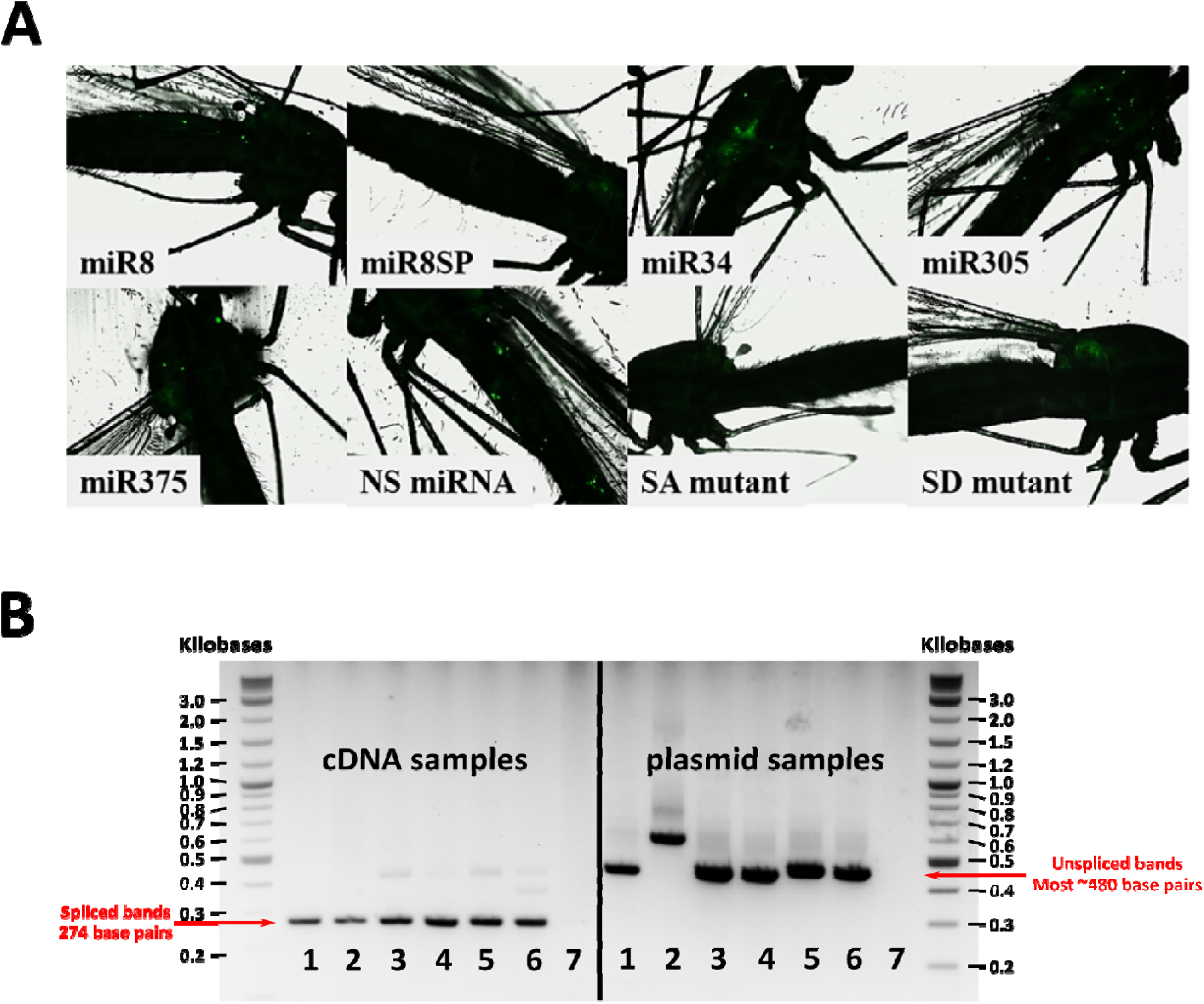
EGFP expression and intronic splicing in 10 days post-injection female mosquitoes. **A)** Punctate expression is visible in mosquitoes co-injected with vWTAgDNV and vAcEGFPmiR8, vAcEGFPmiR8SP, vAcEGFPmiR34, vAcEGFPmiR305, vAcEGFPmiR375, and vAcEGFPNS (panels *miR8*, *miR8SP*, *miR34*, *miR305*, *miR375*, and *NS miRNA*). Little to no EGFP expression was present in mosquitoes co-injected with vWTAgDNV and vAcEGFPSA (*SA* mutant panel). Weak expression is visible in mosquitoes co-injected with vWTAgDNV and vAcEGFPSD (*SD* mutant panel). **B)** PCR product gel measuring intronic splicing in female *An. gambiae* 10 days post-injection with cDNA samples (left side) alongside unspliced DNA plasmid control samples (right side). Matching numbers indicate cDNA and plasmid versions of the same construct. Samples are as follows: 1. *miR8*, 2. *miR8SP*, 3. *miR34*, 4. *miR305*, 5. *miR375*, 6. *Nonsense RNA*, 7. No template control. *In vivo* splicing resulted in a PCR product of 274 bp for wells 1-6 on the cDNA side. A lack of splicing in plasmid controls for wells 1-6 on the plasmid side resulted in bands of ∼480 bp or 572 bp in the case of pAcEGFPmiR8SP.

Similar splicing patterns were observed in Moss55 cells (Fig 3B). Cells co-transfected with pWTAgDNV and transducing plasmid pAcEGFPmiR8, pAcEGFPmiR8SP, pAcEGFPmiR34, pAcEGFPmiR305, pAcEGFPmiR375, or pAcEGFPNS all had 274 bp bands indicative of splicing (cDNA wells 1-6, Fig 3B). Cells co-transfected with pWTDNV and pAcEGFPmiR8, pAcEGFPmiR34, or pAcEGFPNS also had larger ∼480 bp bands consistent with some level of unspliced transcript or splicing intermediates (cDNA wells 1,3, and 6, Fig 3B). In cells co-transfected with pWTAgDNV and pAcEGFPNS, another intermediate-sized band was also present (cDNA well 6, Fig 3B). Cells co-transfected with pWTAgDNV and the splice donor mutant pAcEGFPSD produced a single strong ∼480 bp band representative of a lack of splicing despite faint EGFP expression observed in transfected cells (cDNA well 8, Fig 3B; *SD* mutant, Fig 2B). When cells were co-transfected with pWTAgDNV and the splice acceptor mutant pAcEGFPSA, a ∼480 bp band representative of a lack of splicing as well as a 274 bp band consistent with splicing was observed despite of a lack of EGFP expression in transfected cells (cDNA well 7, Fig 3B; *SA* mutant, Fig 2B). As before, PCRs of plasmid DNA resulted in larger ∼480 bp bands for most constructs (plasmid wells 1 and 3-8, Fig 3B). A band of 572 bp, reflective of a larger intronic segment, was detected for pAcEGFPmiR8SP (plasmid well 2, Fig 3B). Plasmid DNA from pAcEGFP as well as cDNA from cells co-transfected with pWTAgDNV and pAcEGFP lacked the intron and produced bands of 274 bp (plasmid well 10, cDNA well 10, Fig 3B).

Isolations of vWTAgDNV and each transducing virus purified from Moss55 cells were injected into adult female mosquitoes that were 3 days old and images of these mosquitoes were taken 10 days later. Punctate EGFP expression indicative of splicing was observed in the thorax and abdomen of mosquitoes co-injected with vWTAgDNV and vAcEGFPmiR8, vAcEGFPmiR8SP, vAcEGFPmiR34, vAcEGFPmiR305, vAcEGFPmiR375, or vAcEGFPNS (Fig 4A). Little to no EGFP expression was observed in mosquitoes co-injected with vWTAgDNV and vAcEGFPSA (SA mutant, Fig 4A). Mosquitoes injected with vWTAgDNV and vAcEGFPSD exhibited weak EGFP expression that remained localized to the mosquito thorax (SD mutant, Fig 4A). *In vivo* intronic splicing was measured as before via PCR of cDNA made from 10 days post-injection mosquitoes. Spliced 274 bp bands were observed in cDNA samples taken from mosquitoes co-injected with vWTAgDNV and vAcEGFPmiR8, vAcEGFPmiR8SP, vAcEGFPmiR34, vAcEGFPmiR305, vAcEGFPmiR375, or vAcEGFPNS (wells 1-6, cDNA side Fig 4B). Faint unspliced bands were also observed in mosquitoes co-injected with vWTAgDNV and vAcEGFPmiR34, vAcEGFPmiR375, or vAcEGFPNS (cDNA wells 3, 5 and 6, Fig 4B). Plasmid controls resulted in larger bands of ∼480 bp for all constructs except for the larger *miR8SP* insert (plasmid wells 1-6, Fig 4B).

### Preliminary *in vitro* work to select miRNA targets

Preliminary work in Sua5B cells showed that transfection with pWTAgDNV and pAcEGFPmiR8 or pAcEGFPmiR375 led to higher levels of *miR8* and *miR375* respectively, 5 days post-transfection (Fig S1). *miR34* levels were reduced rather than elevated when Sua5B cells were transfected with pWTAgDNV and pAcEGFPmiR34, perhaps indicating processing of this transcript into an anti-miRNA rather than expression of the predicted mature miRNA (Fig S1C). Transfection with pWTAgDNV and pAcEGFPmiR305 or miR8SP did not result in any significant changes in microRNA levels (data not shown). We also observed significant upregulation of the predicted target gene *Cactus* in cells transfected with pWTAgDNV and pAcEGFPmiR375 (Fig S2). Given the lack of significant change in miRNA levels for *miR8SP*, and *miR305*, and difficulty in interpreting results from *miR8* in light of *miR8SP* results, these miRNAs were not evaluated in later experiments. A preliminary *in vivo* experiment was also carried out where mosquitoes injected with vWTAgDNV and vAcEGFPmiR34 showed significant elevation in *miR34* target gene transcripts *MISO*, *Caspar*, and *Rel2* although the level of *miR34* was not changed in mosquitoes at this timepoint (Fig S3).

### *In vivo* infections

After down-select, larger-scale *in vivo* experiments focused on mosquitoes injected with vWTAgDNV and either vAcEGPFmiR34 or vAcEGFPmiR375. Mosquitoes injected with vWTAgDNV and vAcEGPFmiR34 were evaluated for changes in *MISO*, *Caspar*, and *Rel2* transcript levels whereas mosquitoes injected with vWTAgDNV and vAcEGPFmiR375 were tested for differences in *Cactus* and *Rel1A* transcript levels.

*mir34* levels were not significantly different between mosquitoes injected with vWTAgDNV and vAcEGFPmiR34 and control mosquitoes injected with vWTAgDNV and vAcEGFPNS (Fig 5A), similar to preliminary *in vivo* results. No differences were seen in *Caspar* or *MISO* transcript levels 10 days post-injection (Fig 5B and 5C). However, *Rel2* transcripts were more abundant 10 days post-infection in mosquitoes injected with vWTAgDNV and vAcEGFPmiR34 compared to those injected with vWTAgDNV and vAcEGFPNS (Fig 5D). The direction of this change in target gene transcript abundance may indicate miRNA signaling through the lessor known RNA activation pathway rather than the RNAi pathway or that this miRNA acts on an upstream inhibitor of *Rel2* [60–62].

**Figure 5:**
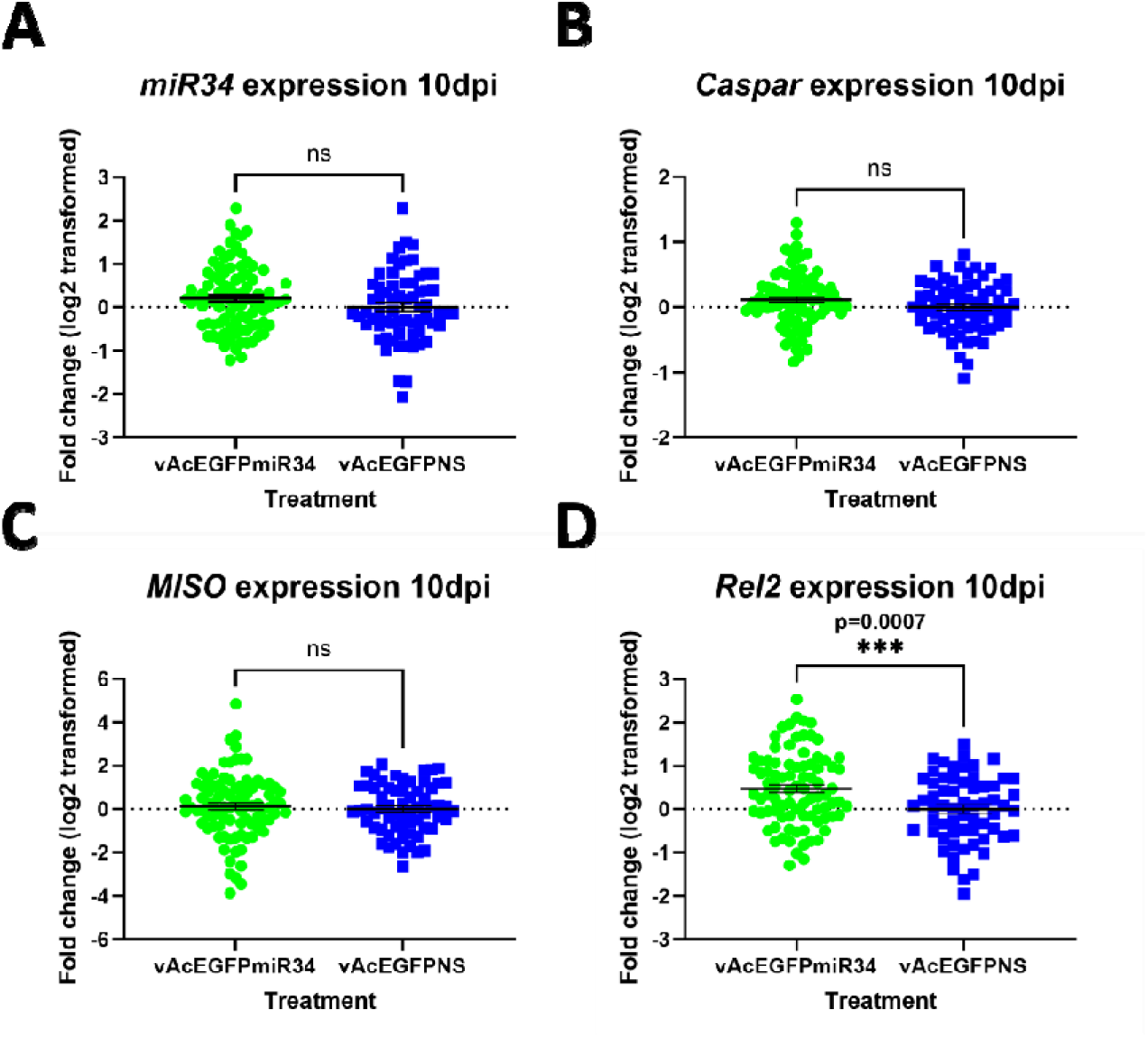
Ex pression o f *miR34*, and target gene transcripts in mosquitoes 10 days post-injection with vWTAgDNV and vAcEGFPmiR34. **A)** Expression of *miR34* was unchanged in mosquitoes that were injected with vWTAgDNV and vAcEGFPmiR34. **B)** Expression of *Caspar* transcripts was not altered following injection. **C)** Levels of *MISO* were also not changed 10 days post-injection. d. *MISO* transcript levels were significantly elevated following injection. D) *Rel2* expression was enhanced in mosquitoes following injection of vWTAgDNV and vAcEGFPmiR34. Dashed lines indicate a fold change of 0. Green dots represent individual mosquitoes co-injected with vAcEGFPmiR34 and pWTAgDNV whereas blue squares indicate individual mosquitoes co-injected with control vAcEGFPNS and vWTAgDNV. Data in A-D were normal as assessed by a D’Agostino-Pearson normality test and were analyzed using an unpaired t-test.

When mosquitoes were injected with vWTAgDNV and vAcEGFPmiR375, *miR375* levels were slightly reduced 10 days post-infection (Fig 6A). This was unexpected given the strong increases seen in preliminary *in vitro* experiments and may indicate that the introduced miRNA is not processed *in vivo* as expected and instead produces an anti-miRNA that binds to endogenous *miR375*. Both *Cactus* and *Rel1A* transcript levels were slightly elevated in mosquitoes 10 days post-infection (Fig 6B and 6C). This contrasts with the predicted increase in *Cactus* and decrease in *Rel1A* from *Ae. aegypti*.

**Figure 6:**
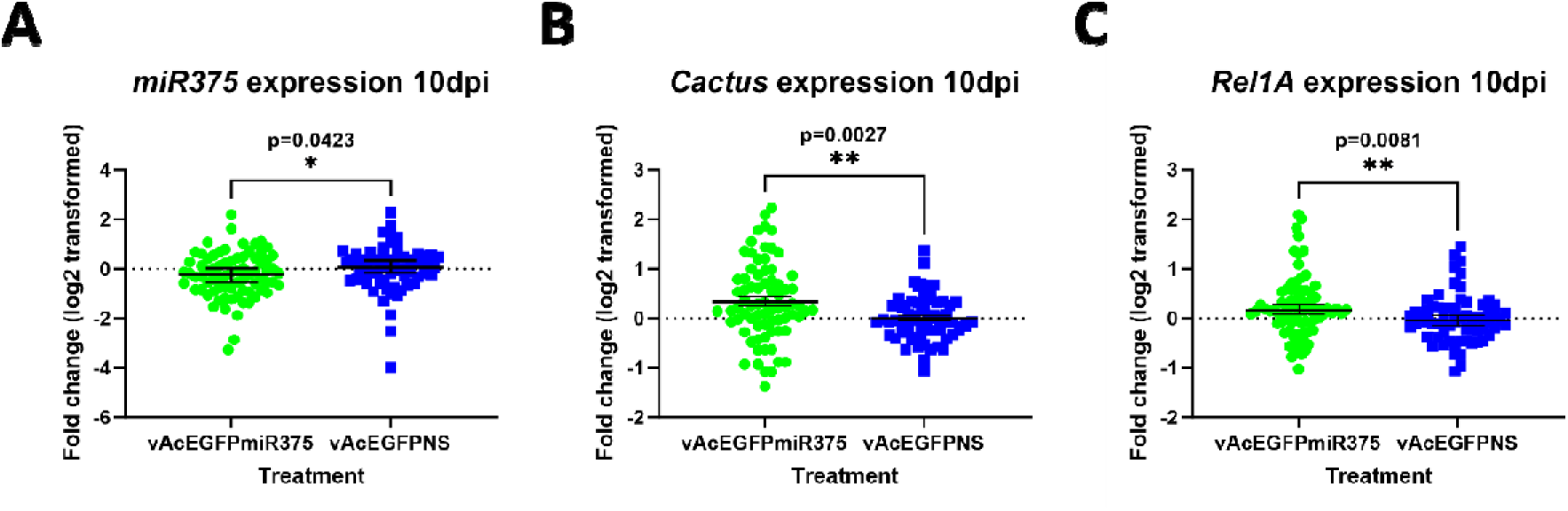
Expression of *miR375*, and target gene transcripts in mosquitoes 10 days post-injection with vWTAgDNV and vAcEGFPmiR375. **A)** Expression of *miR375* was slightly decreased in mosquitoes injected with vWTAgDNV and vAcEGFPmiR375. **B)** Expression of *Cactus* transcripts was elevated in mosquitoes 10 days post-injection. **C)** Levels of *Rel1A* expression was slightly increased 10 days post-injection with vWTAgDNV and vAcEGFPmiR375. Dashed lines indicate a fold change of 0. Green dots represent individual mosquitoes co-injected with vAcEGFPmiR375 and pWTAgDNV whereas blue squares indicate individual mosquitoes co-injected with control vAcEGFPNS and vWTAgDNV. Data in A and B was not normally distributed as assessed by a D’Agostino-Pearson normality test and was analyzed using a nonparametric Mann-Whitney test. Data in C was normally distributed and analyzed using an unpaired t-test.

## Discussion

These experiments show that splicing of the developed AgDNV-delivered intron is robust both *in vitro* within two different *An. gambiae* cell lines of varied lineages, as well as *in vivo* for a variety of endogenous pre-miRNA sequences, one developed miRNA sponge sequence, and one random RNA sequence. Occasionally, unexpected unspliced transcripts were observed in PCR assays. These likely represent splicing intermediates or pre-splicing transcripts as the presence of these larger bands did not correlate with altered EGFP expression *in vitro* or *in vivo* (Fig 2A and 2B, Fig 4A). This demonstrates that splicing is specific to the developed intronic sequence and not altered by the cargo sequence. Further supporting this, intronic splicing can be eliminated or reduced through alteration of *splice donor* or *splice acceptor* sequences. Although mutated *splice donor* and *splice acceptor* constructs within the two cell lines sometimes produced bands seemingly consistent with some level of splicing, reduced or absent *in vitro* EGFP expression observed for these constructs indicates that any splicing that occurs is greatly suppressed, modified, or results a stop codon as predicted (cDNA wells 7 and 8, Fig 3A, cDNA well 7, Fig 3B, SA and SD mutants Fig 2A-B). Although splicing was not assessed by PCR *in vivo* for *splice donor* and *splice acceptor* mutants, mosquitoes injected with vWTAgDNV and vAcEGFPSA had little to no EGFP expression and mosquitoes injected with vWTAgDNV and vAcEGFPSD exhibited weak EGFP expression (SA and SD mutants, Fig 4A). Thus, this intron and expression strategy represents a promising new method for introducing small RNAs both *in vitro* and *in vivo*.

Despite the success of this expression strategy *in vitro*, some differences between cell types were observed. Sua5B cells are larger in size and produced noticeably stronger EGFP expression than Moss55 cells (Fig 2A and 2B). While not investigated in this study, this difference in observed EGFP expression may be due to chronic AgDNV infection already present in Sua5B cells that could enhance viral replication and packaging efficiency of transducing constructs [13]. Moss55 cells lack natural AgDNV infection and thus rely solely on AgDNV constructs expressed from transfected plasmids. Alternatively, Sua5B cells are considered to have hemocyte-like properties whereas Moss55 cells have an epithelial origin and differences in cell lifecycle or rate of transcription may explain the variation in EGFP intensity [63–65]. Despite these differences in AgDNV infection status and EGFP intensity, no differences were observed in the viral titers produced by the different cell lines for mosquito injections and to better control the genetic diversity of purified viruses, Moss55 cells were used to grow viral stocks used for *in vivo* experiments.

*In vivo* experiments largely focused on mosquitoes 10 days post-injection with vWTAgDNV and either vAcEGFPmiR34 or vAcEGFPmiR375. While experiments with mosquitoes injected with vWTAgDNV and vAcEGFPmiR34 did not result in any differences in *mir34* levels, *Rel2* transcript levels were slightly elevated 10 days post-infection (Fig 5A and 5D). Although a directionality to *Rel2* changes was not predicted prior to experiments, this increase in *Rel2* transcript abundance indicates that *miR34* may act on *Rel2* through the RNA activation pathway or that *miR34* acts on another transcript that in turn influences *Rel2* abundance, possibly an upstream inhibitor of *Rel2* (Table 1) [60–62]. Similarly, although *miR375* was predicted to bind to *Cactus* and *Rel1* and cause an increase in *Cactus* and decrease in *Rel1*, mosquitoes that were injected with vWTAgDNV and vAcEGFPmiR375 had slightly elevated *Cactus* and *Rel1A* transcript levels 10 days post-infection (Fig 6B and 6C). This was unexpected as *Cactus* is a negative regulator of *Rel1* and if binding of *miR375* results in transcript depletion though RNAi, in theory, binding would decrease the transcript levels of both. As with *miR34*, it is possible that *miR375* is acting through RNAa or on transcripts upstream of *Cactus* or *Rel1*. Additionally, as *miR375* levels were slightly decreased in mosquitoes infected with vWTAgDNV and vAcEGFPmiR375, it is also possible that this miRNA is being processed into an anti-miRNA and that this binds to mature *miR375* or operates in a different way than mature *miR375* (Fig 6A).

There are many possible reasons for differences between what we observed from *in vivo* expression experiments versus what was expected based on prior predictions (Table 1). Most importantly, although there have been some prior studies for certain miRNAs, the true direct targets of these *An. gambiae* miRNAs have largely not been identified and most targets have only been bioinformatically predicted (Table 1). Further, muted changes in miRNA or target gene transcripts could be due to organ specific effects that are diluted when analyzing the whole mosquito body as we did. Inconsistencies could also be due to a lack of effective miRNA expression *in vivo* or indicate that infection levels of transducing viruses were lower than those needed to induce a significant alteration in endogenous miRNAs. Despite these inconsistencies between predicted and observed alterations, we did observe changes in *miR375* levels and in some *miR34* and *miR375* target transcripts indicating that the developed artificial intron has potential for use in miRNA expression and may be more successful if specific target genes are better characterized. (Fig 5D, Fig 6A-C).

Future experiments should examine miRNA and target gene transcript levels within organs known to support AgDNV infection such as the midgut, ovaries, and fat body. By examining specific organs rather than the whole mosquito, noise in the system may be reduced and changes in miRNAs or target genes may be more evident. The system can also such as conceptually be useful for expressing other effectors such as small interfering RNAs (siRNAs) developed against specific genes. By expressing siRNAs with specific targets, the true capabilities of this virus and intron expression system could be better tested and assessed for meaningful intervention in biological or field settings. Additional studies of AgDNV viral replication dynamics, dosing requirements for injections, and organ specificity would greatly aid future work and the further development of this symbiont as a tool for paratransgenesis.

## Acknowledgements

This research was supported by NIH/NIAID grant R01AI12820, NSF/BIO grant 1645331, USDA Hatch Project 4769, a grant with the Pennsylvania Department of Health using Tobacco Settlement Funds, and funds from the Dorothy Foehr Huck and J. Lloyd Huck endowment to JLR.

## Supplemental data

**Table S1:**
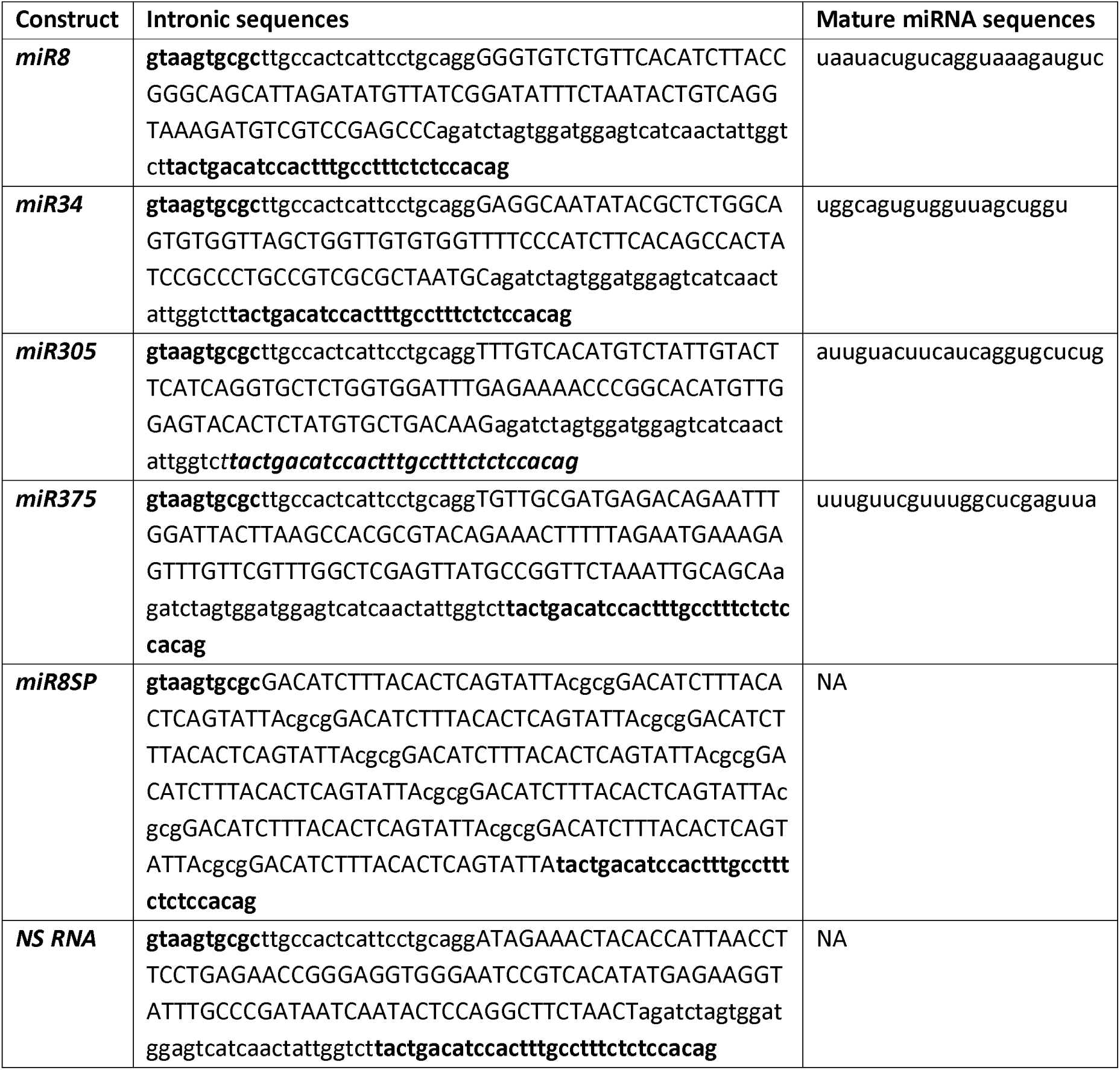
Intronic sequences for transducing constructs. Bold font indicates the intronic segments of the splice donor and splice acceptor site. Uppercase font indicates the pre-miRNA sequences for miRNAs (*miR8*, *miR34*, *miR305*, or *miR375*), or the miRNA sponge blocks that are the reverse complement of *miR8* (*miR8SP*), or the *nonsense RNA* sequence (*NS RNA*). Lowercase, non-bold font indicates spacer sequences. Mature miRNA sequences are given for *miR8*, *miR34*, *miR305*, and *miR375*.

**Table S2:**
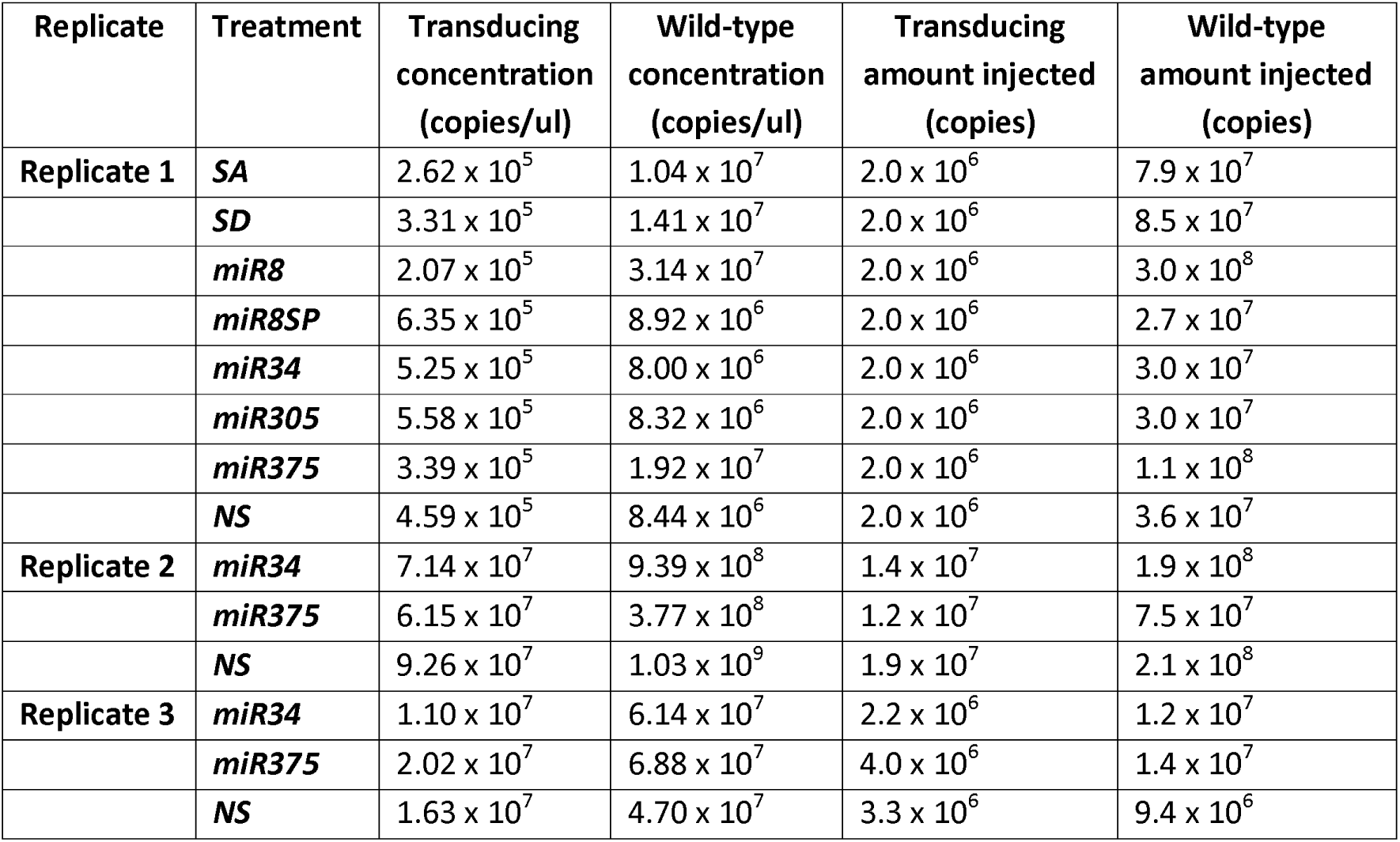
Concentrations of transducing and WT virus used to inject mosquitoes. Listed values in copies/ul were found using qPCR against WT AgDNV *nonstructural protein 1* for vWTAgDNV and a standard curve of pWTAgDNV. For transducing virus, qPCR reactions against *EGFP* and standard curve of pAcEGFP was used to calculate copies/ul. Replicate 1 samples were concentrated to ∼1 × 10^7^ copies/ul of the transducing virus using an Amicon Ultra 0.5 mL Ultracel 10K filters (UFC501096) before injection. Injection values for Replicate 1 reflect the concentrated amount injected whereas the transducing and wild-type concentration represent the copies/ul measured by qPCR before concentrating. Injected copies were calculated using an injection volume of 200 nl per mosquito. Replicate 2 and Replicate 3 viruses did not need to be concentrated before injection.

**Table S3:**
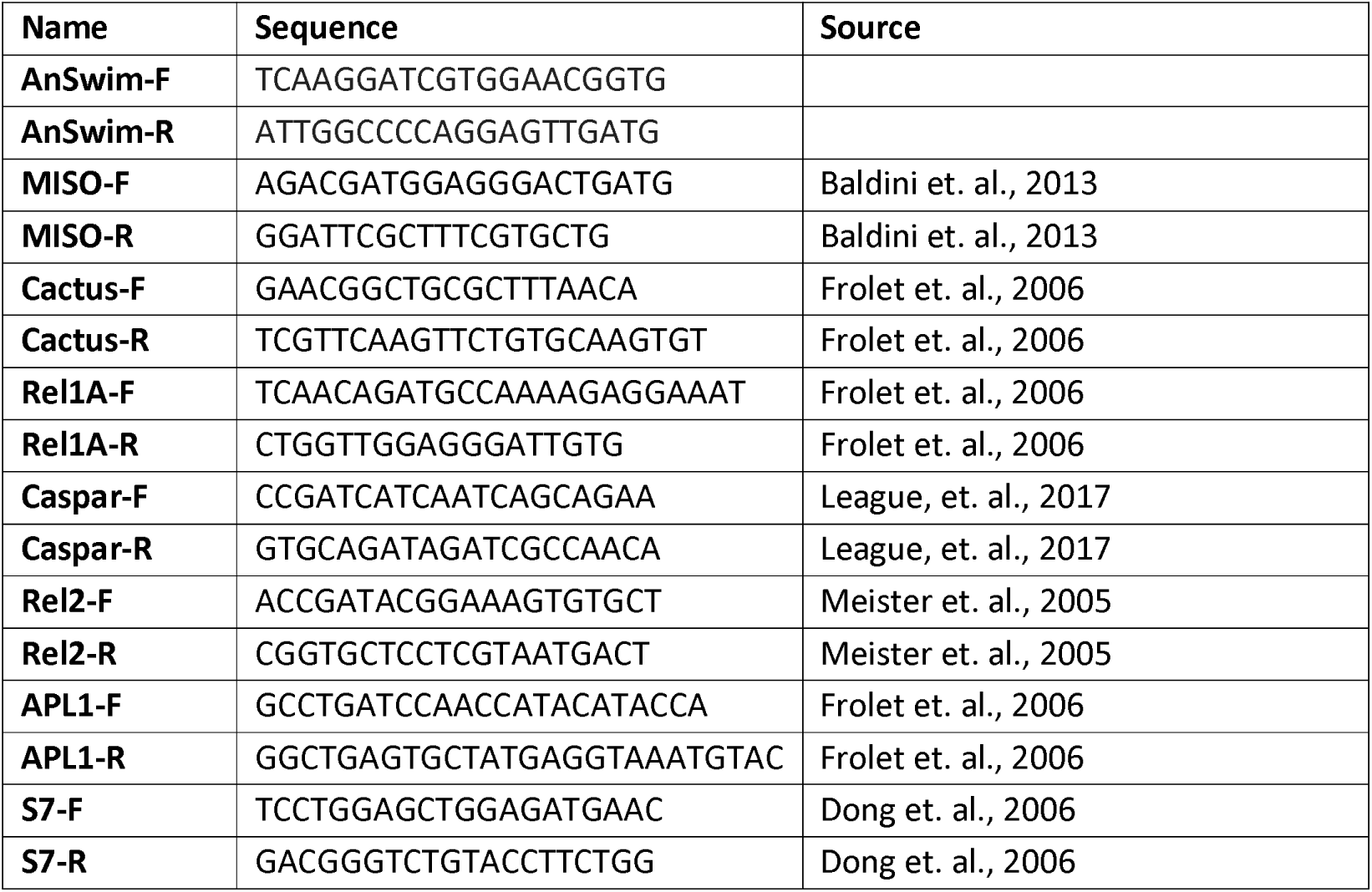
Primers used for target gene qPCR reactions. Primers were taken from the reported sources except for primers targeting *Swim1* that were developed during this study.

**Table S4:**
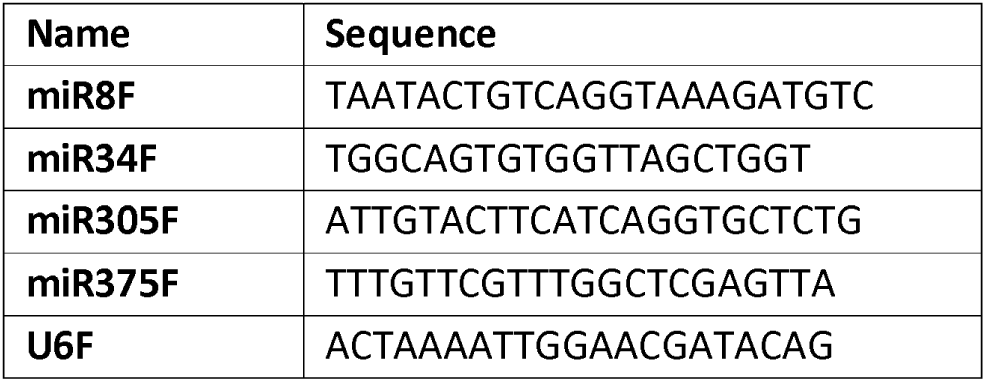
Primers used for miRNA quantification qPCR reactions. Primers matching the sequence of each miRNA were developed against each mature miRNA sequence. *U6* primer was developed against *An. gambiae U6* and this small RNA served as a reference with which to compare miRNA expression.

**Figure S1:**
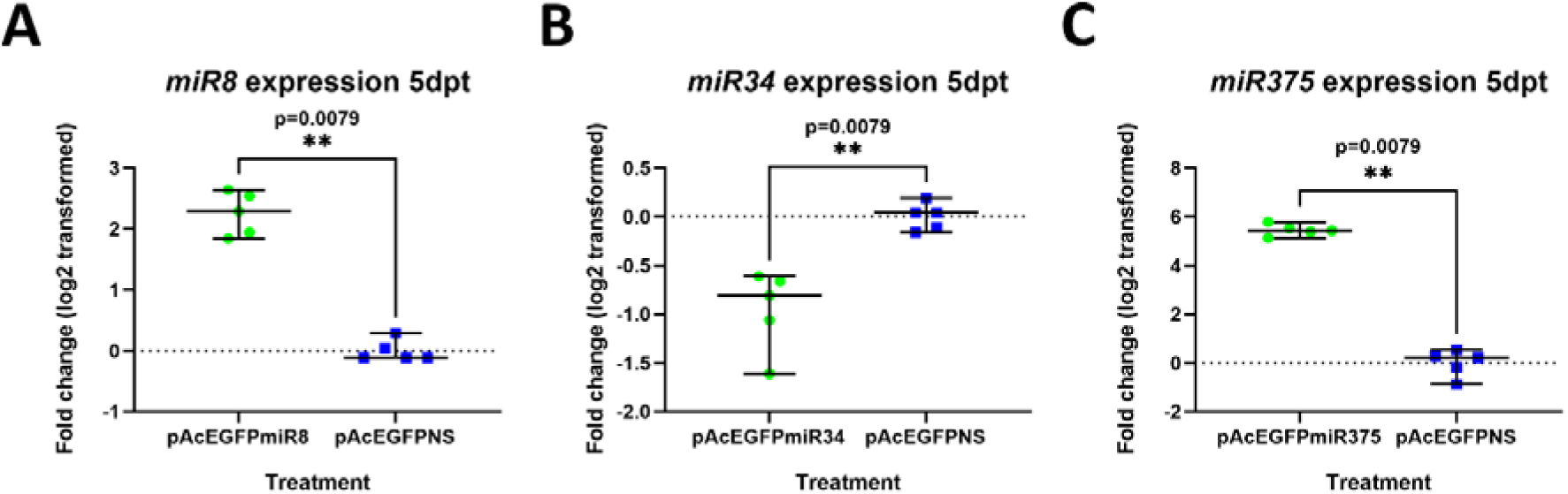
Expression of *miR8*, *miR34*, and *miR375* in Sua5B cells 5 days post-transfection. **A)** Levels of *miR8* were elevated when cells were co-transfected with pWTAgDNV and pAcEGFPmiR8. **B)** Levels of *miR34* were lower when cells were transfected with pWTAgDNV and pAcEGFPmiR34. **C)** miR375 expression was increased when cells were transfected with pWTAgDNV and pAcEGFPmiR375. Dashed line indicates a fold change of 0. Green dots represent individual wells of cells co-transfected with indicated miRNA-expressing plasmids and pWTAgDNV whereas blue squares indicate individual wells co-transfected with control pAcEGFPNS and pWTAgDNV. Data in all graphs did not follow a normal distribution and failed D’Agostino-Pearson normality tests. Differences between groups were analyzed using Mann-Whitney tests.

**Figure S2:**
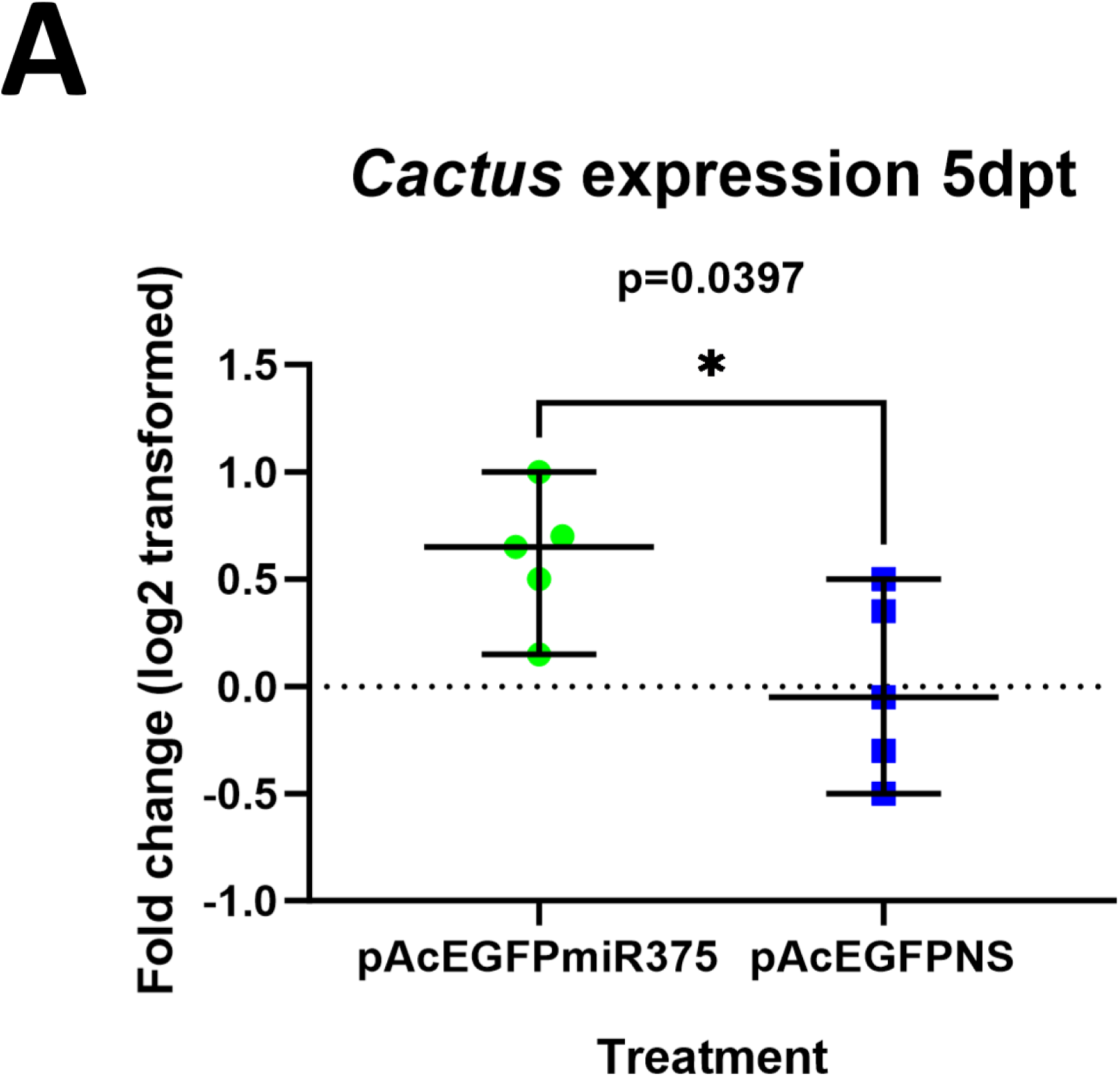
Expression of *Cactus* in Sua5B cells 5 days post-transfection with pAcEGFPmiR375 and pAcEGFPNS. **A)** *Cactus* transcript levels were elevated when cells were co-transfected with pWTAgDNV and pAcEGFPmiR375. Dashed line indicates a fold change of 0. Green dots represent individual wells of cells co-transfected with pAcEGFPmiR375 and pWTAgDNV whereas blue squares indicate individual wells co-transfected with control pAcEGFPNS and pWTAgDNV. Data and failed a D’Agostino-Pearson normality test and was analyzed using a Mann-Whitney test.

**Figure S3:**
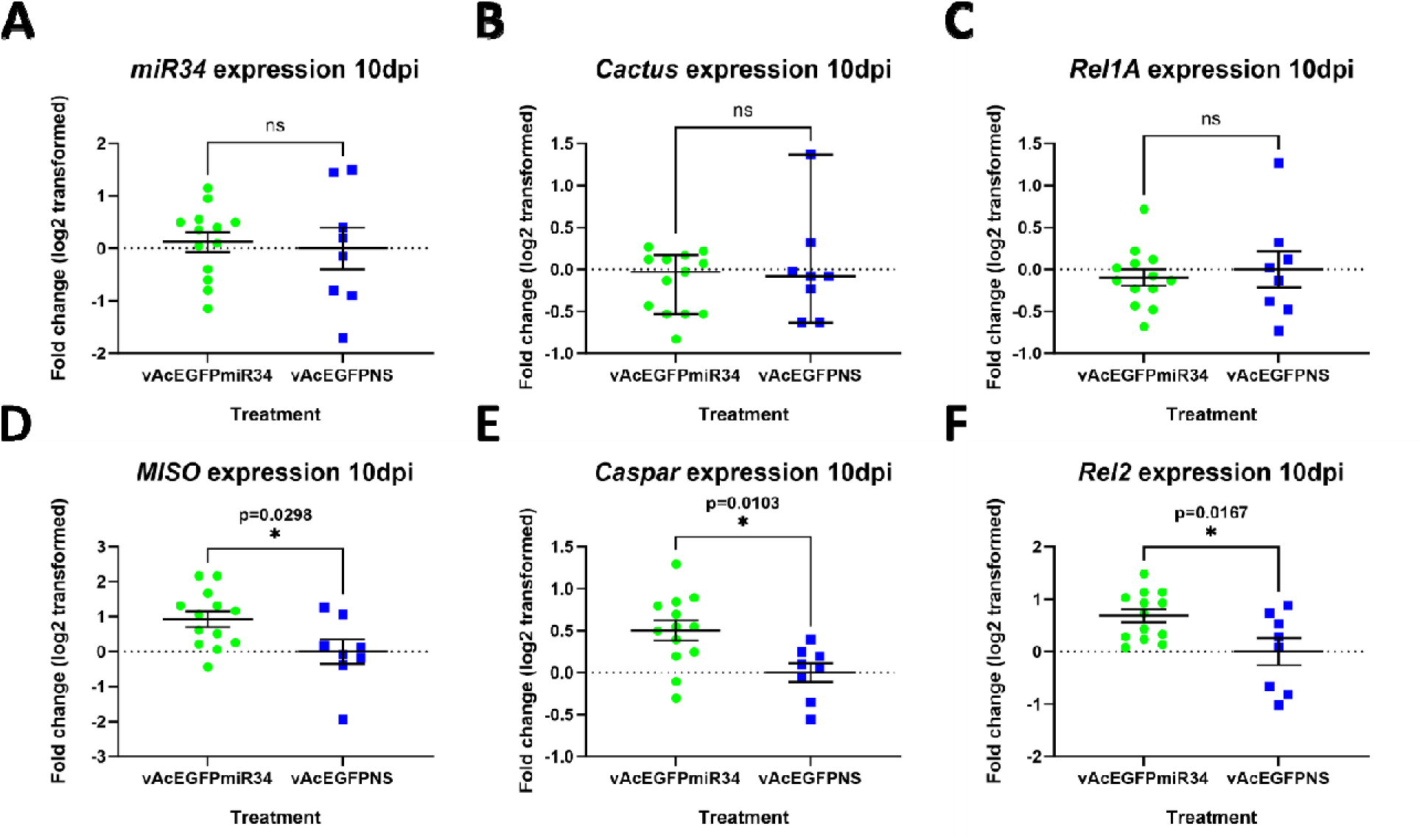
Expression of *miR34*, and target gene transcripts in replicate 1 mosquitoes 10 days post-injection with vWTAgDNV and vAcEGFPmiR34. **A)** Expression of *miR34* was unchanged in mosquitoes that were injected with vWTAgDNV and vAcEGFPmiR34. **B)** *Cactus* transcript levels were not altered following injection. **C)** Levels of *Rel1A* were also not changed 10 days post-injection. **D)** *MISO* levels were significantly elevated following injection. **E)** *Caspar* expression was enhanced in mosquitoes following injection of vWTAgDNV and vAcEGFPmiR34. **F)** *Rel2* expression was also enhanced in mosquitoes following injection. Dashed line indicates a fold change of 0. Green dots represent individual mosquitoes co-injected with vAcEGFPmiR34 and pWTAgDNV whereas blue squares indicate individual mosquitoes co-injected with control vAcEGFPNS and vWTAgDNV. Data in graphs A and C-F were normal as assessed by a D’Agostino-Pearson normality test and were analyzed using an unpaired t-test. Data in graph B were not normal and were analyzed using a Mann-Whitney test.

